## Supplementary Information - Figures and Tables for "Multiple T6SSs, mobile auxiliary modules, and effectors revealed in a systematic analysis of the *Vibrio parahaemolyticus* pan-genome"

**Supplementary Figures S1-S6**

**Supplementary Tables S1-S2**

**Supplementary Dataset S1-S6**

**Supplementary Movies S1**

**Supplementary References**

### Supplementary Figures

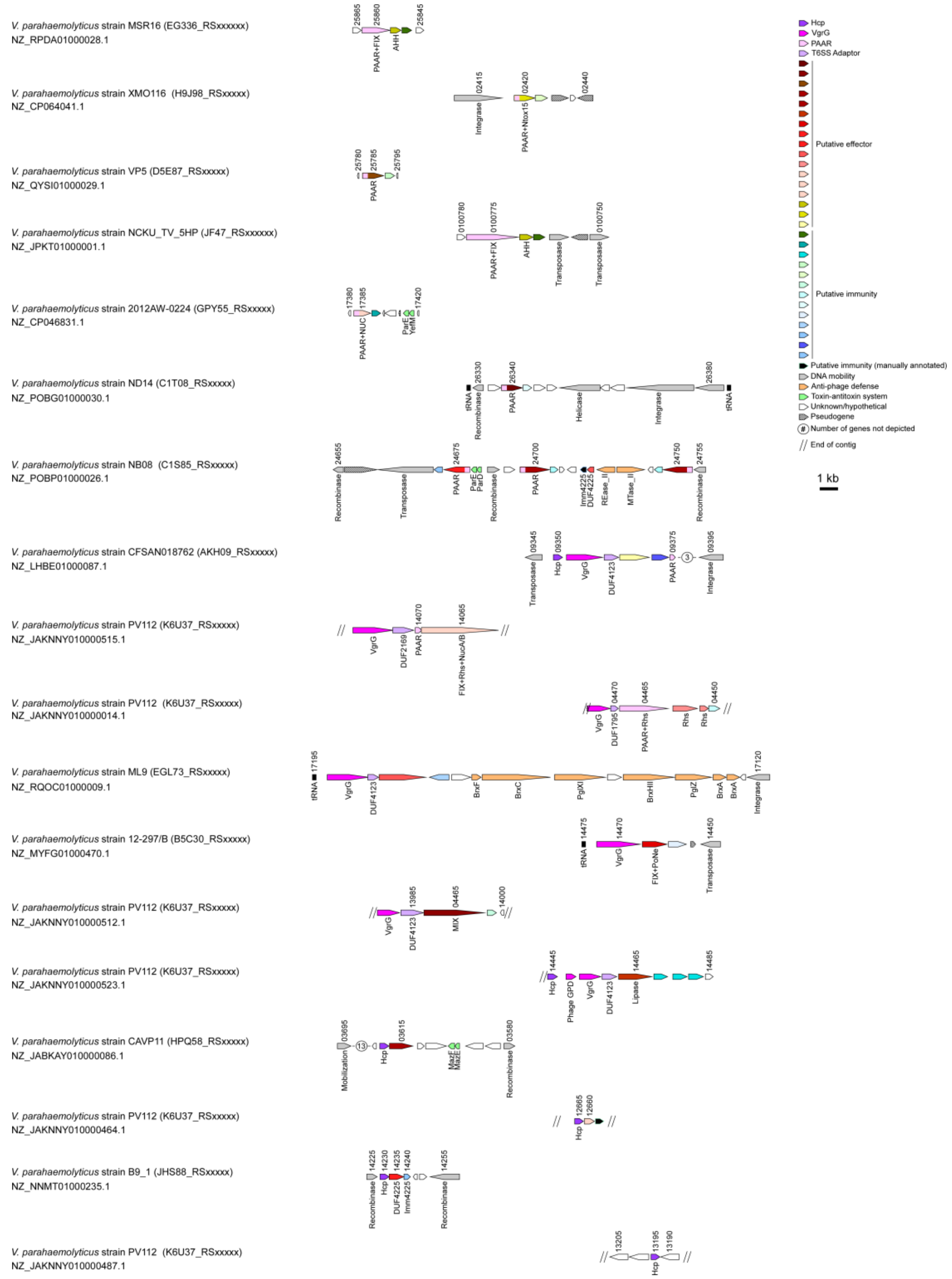


**Supplementary Figure S1. T6SS auxiliary module types in *V. parahaemolyticus*.** Representative T6SS auxiliary modules found in *V. parahaemolyticus* genomes. The strain names, the GenBank accession numbers, and the locus tag annotation patterns are provided. Genes are denoted by arrows indicating the direction of transcription. Locus tags are denoted above, and the names of encoded proteins or domains are denoted below.

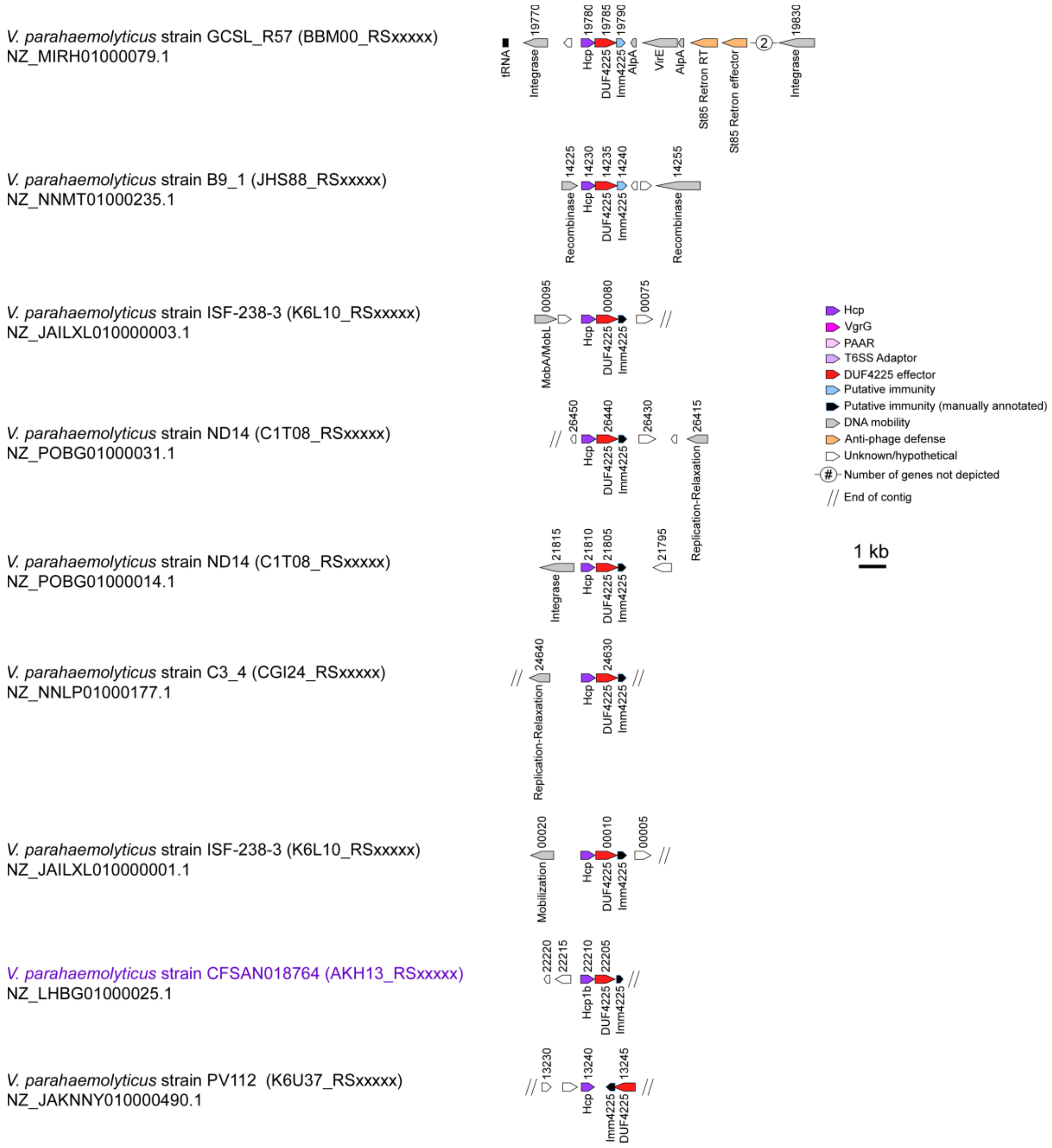


**Supplementary Figure S2. DUF4225-encoding Hcp T6SS auxiliary module types in *V. parahaemolyticus.*** Representative Hcp auxiliary modules containing a DUF4225-encoding gene found in *V. parahaemolyticus* genomes. The strain names, the GenBank accession numbers, and the locus tag annotation patterns are provided. Genes are denoted by arrows indicating the direction of transcription. Locus tags are denoted above, and the names of encoded proteins or domains are denoted below. The Hcp1b module investigated in this work is denoted in purple.

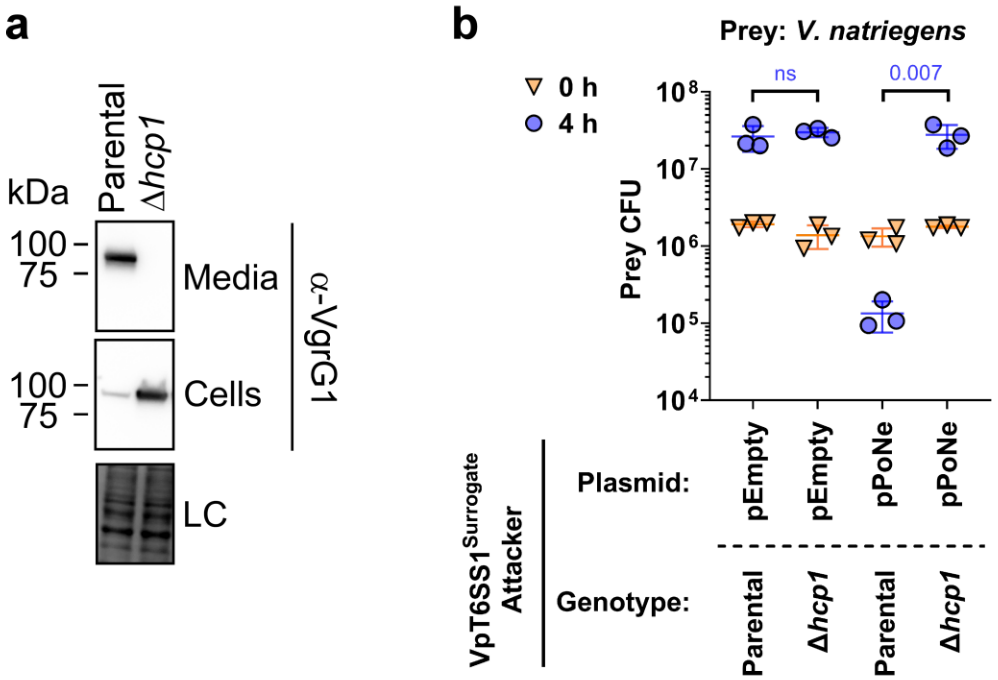


**Supplementary Figure S3. Constructing a functional effectorless T6SS1 surrogate platform. (a)** Expression (cells) and secretion (media) of VgrG1 from the indicated VpT6SS1$^{Surrogate}$ strain or its Δ*hcp1* derivative. Samples were grown in MLB media at 30°C. Loading control (LC), visualized as trihalo compounds' fluorescence of the immunoblot membrane, is shown for total protein lysates. **(b)** Viability counts (CFU) of *V. natriegens* prey strain before (0 h) and after (4 h) co-incubation with the surrogate T6SS1 platform strain (VpT6SS1$^{Surrogate}$) or its T6SS1$^{-}$ derivative (Δ*hcp1*) carrying an empty plasmid (pEmpty) or a plasmid for the arabinose-inducible expression of the PoNe DNase-containing VgrG1b module from *V. parahaemolyticus* 12-297/B (pPoNe). The statistical significance between samples at the 4 h timepoint was calculated using an unpaired, two-tailed Student's *t*-test; ns, no significant difference ($p > 0.05$). Data are shown as the mean ± SD; n = 3.

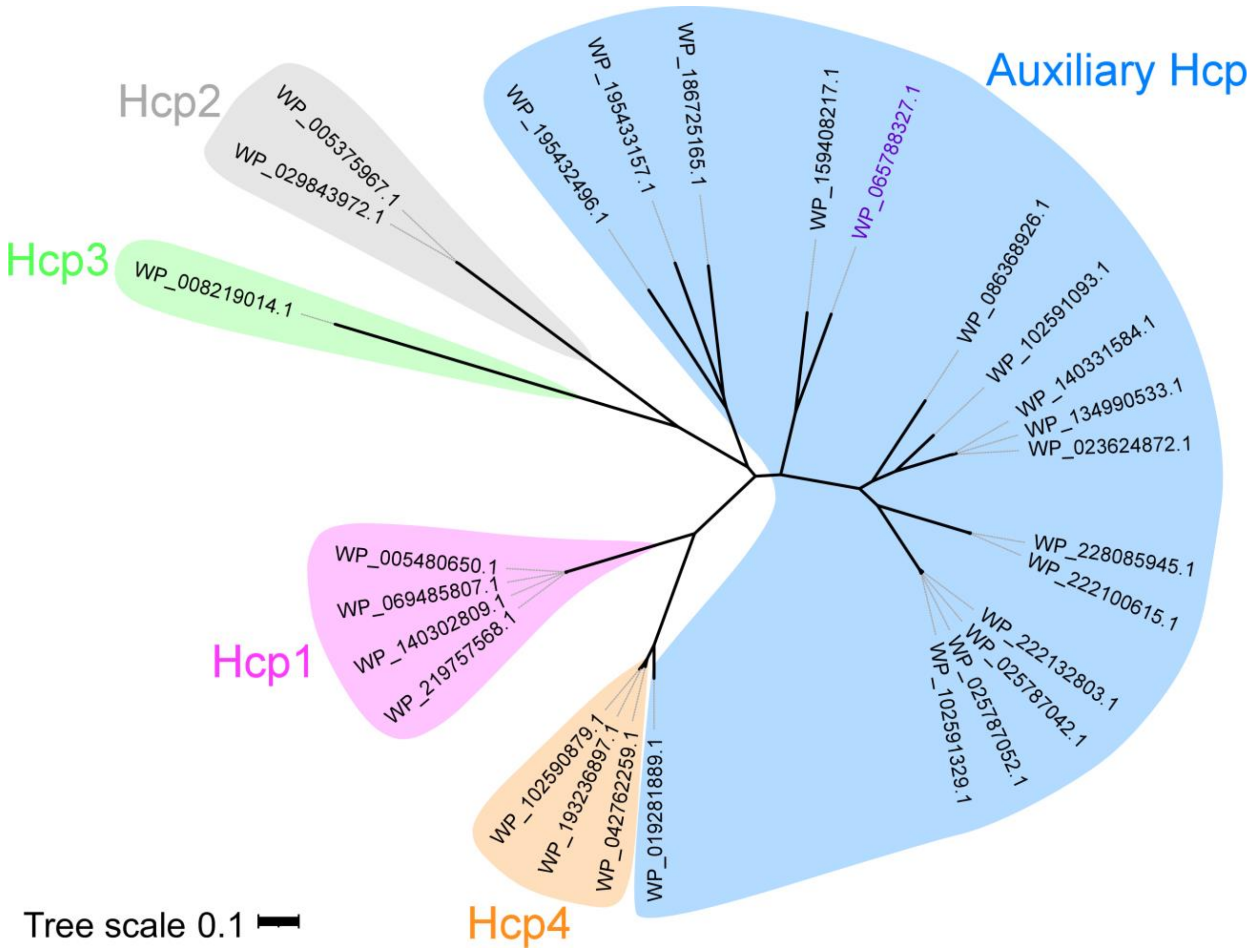


**Supplementary Figure S4. Phylogenetic distribution of Hcp encoded within *V. parahaemolyticus* genomes.** The evolutionary history was inferred using the neighbor-joining method. The phylogenetic tree was drawn to scale, with branch lengths in the same units as those of the evolutionary distances used to infer the phylogenetic tree. The evolutionary distances were computed using the Poisson correction method and are in the units of the number of amino acid substitutions per site. The protein accession numbers are denoted. The Hcp1b investigated in this work is denoted in purple.

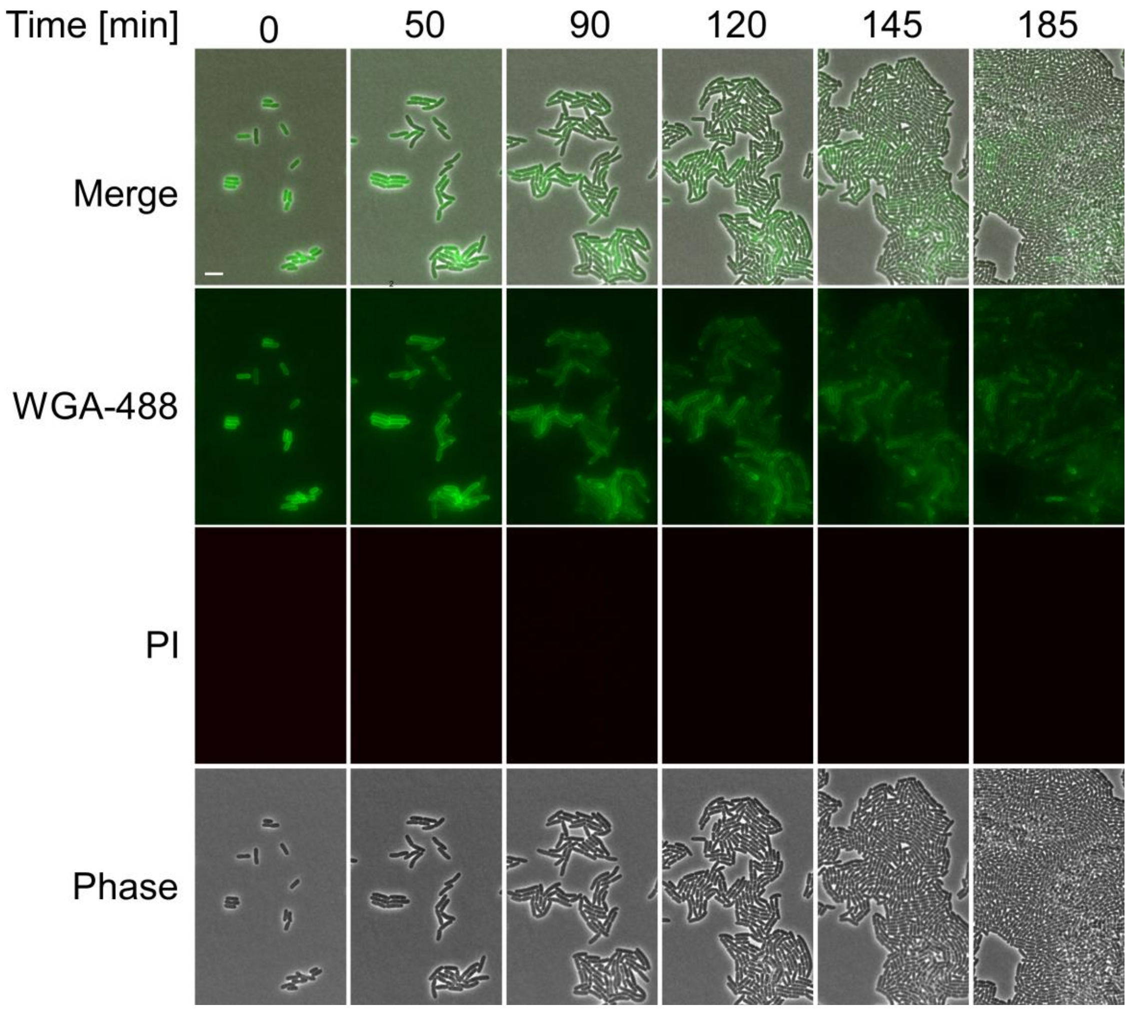


**Supplementary Figure S5. Time-lapse microscopy of *E. coli* cells.** *E. coli* MG1655 cells stained with Wheat Germ Agglutinin Alexa Fluor 488 conjugate (WGA-488) and that contain an empty arabinose-inducible expression plasmid, grown on agarose pads supplemented with chloramphenicol (to maintain the plasmid) and 0.2% arabinose (to induce expression), and propidium iodide (PI). WGA-488 (green), PI (red), phase contrast and merged channels are shown. Size bar = 5 µm.

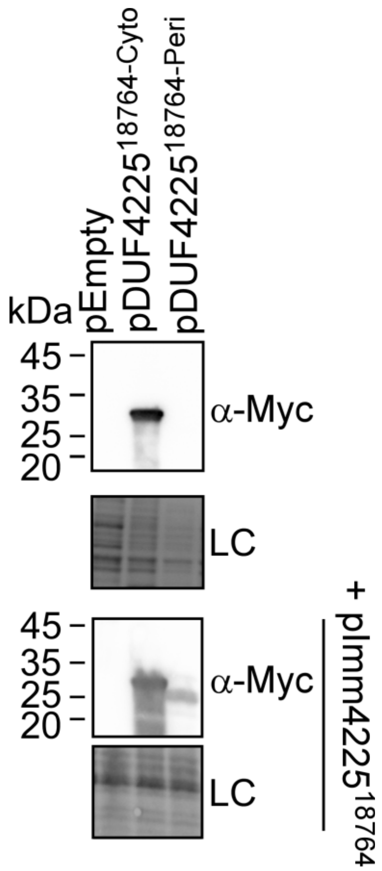


**Supplementary Figure S6. Expression of DUF4225$^{18764}$ in *E. coli*.** Expression of the indicated C-terminal Myc-His-tagged DUF4225$^{18764}$ variants from pBAD$^{K}$/Myc-His (cytoplasmic; Cyto) or pPER5 (periplasmic; Peri) in *E. coli* BL21 (DE3), in the absence (top panels) or presence (bottom panels) of Imm4225$^{18764}$ expressed from pBAD33.1$^{F}$ (pImm4225$^{18764}$). DUF4225$^{18764}$ proteins were detected by immunoblotting using specific α-Myc antibodies. Loading control (LC), visualized as trihalo compounds' fluorescence of the immunoblot membrane, is shown for total protein lysates.

### Supplementary Tables

#### Supplementary Table S1. A list of bacterial strains used in this work.

| Strain name | Genotype | Comments | Source |
|---|---|---|---|
| *Vibrio parahaemolyticus* RIMD 2210633 | Wild type | Used for generating deletion strains. | (1) |
| *Vibrio parahaemolyticus* RIMD 2210633 Δ*hcp1* | Δ*vp1393* | RIMD 2210633 derivative containing an in-frame deletion of *vp1393;* used as prey in bacterial competition assays. | (2) |
| VpT6SS1$^{Surrogate}$ | Δ*tdhAS*/Δ*vp1388/*Δ*vpa1263/ vp1415$^{AAA}$*/Δ*vp1133* | RIMD 2210633 derivative containing in-frame deletions of *tdhAS*, *vp1388*, *vpa1263*, and *vp1133*, and substitutions of the codons encoding histidines 563-4 in VP1415 with codons for alanines. Used as a surrogate platform in competition assays and in secretion assays. | This study |
| VpT6SS1$^{Surrogate/\Delta hcp1}$ | Δ*tdhAS*/Δ*vp1388/*Δ*vpa1263/ vp1415$^{AAA}$*/Δ*vp1133/* Δ*vp1393* | VpT6SS1$^{Surrogate}$ derivative containing an in-frame deletion of *vp1393;* used as a T6SS1$^{-}$ surrogate platform in competition assays and in secretion assays. | This study |
| *Vibrio natriegens* ATCC 14048 | Wild type | Used as prey in competition assays. | ATCC collection |
| *Vibrio coralliilyticus* ATCC BAA-450 | Wild type | Used for generating deletion strains. | ATCC collection |
| *Vibrio coralliilyticus* ATCC BAA-450 Δ*hcp1* | Δ*vic_rs16330* | ATCC BAA-450 derivative containing an in-frame deletion of *vic_rs16330*; used as prey in competition assays. | This study |
| *Vibrio campbellii* ATCC 25920 | Wild type | Used for generating deletion strains. | ATCC collection |
| *Vibrio campbellii* ATCC 25920 Δ*hcp1* | Δ*a8140_16775* | ATCC 25920 derivative containing an in-frame deletion of *a8140_16775;* used as prey in competition assays. | This study |
| *Vibrio parahaemolyticus* 12-297/B Δ*hcp1* | Δ*b5c30_rs15290* | 12-297/B derivative containing an in-frame deletion of *b5c30_rs15290;* used as prey in competition assays | (3) |
| *Vibrio vulnificus* CMCP6 | Rifampicin-resistant | Used as prey in competition | Obtained |

| | | | |
|---|---|---|---|
| | parental strain | assays. | from Karla Satchell |
| *Aeromonas jandaei* DSM 7311 Δ*tssB* | Δ*bn1126_rs13720* | DSM 7311 derivative containing an in-frame deletion of *bn1126*_rs13720; used as prey in competition assays | (4) |
| *Escherichia coli* DH5α (λ pir) | K-12 derivative laboratory strain containing λ pir | Used for plasmid maintenance and cloning. | Obtained from Eric V. Stabb |
| *Escherichia coli* BL21 (DE3) | Laboratory strain | Used for protein expression and toxicity assays. | Obtained from Kim Orth |
| *Escherichia coli* MG1655 | Laboratory strain | Used for microscopy assays. | |

**Supplementary Table S2. A list of plasmids used in this work.**

| Plasmid name | Description | Purpose | Source |
|---|---|---|---|
| $pBAD^{K}$/Myc-His | pBR322 ori-containing plasmid harboring a $Kan^{R}$ cassette, *araC*, and an MCS following a P*bad* promoter | Used for arabinose-inducible expression | (5) |
| $pDUF4225^{18764\text{-}Cyto}$ | $pBAD^{K}$/Myc-His plasmid containing the CDS of $DUF4225^{18764}$ in frame with the C-terminal Myc-His tag | Used for arabinose-inducible expression of $DUF4225^{18764}$ proteins in *E. coli* | This study |
| pPER5 | $pBAD^{K}$/Myc-His with a PelB signal peptide inserted at the 5' end of the MCS | Used for arabinose-inducible expression of proteins targeted to the periplasm in *E. coli* | (6) |
| $pDUF4225^{18764\text{-}Peri}$ | pPER5 plasmid containing the CDS of $DUF4225^{18764}$ in frame with the N-terminal PelB signal peptide and the C-terminal Myc-His tag | Used for arabinose-inducible expression of $DUF4225^{18764}$ protein targeted to the periplasm in *E. coli* | This study |
| $pTme1^{peri}$ | pPER5 plasmid containing the CDS of Tme1 from *V. parahaemolyticus* BB22OP in frame with the N-terminal PelB signal peptide and the C-terminal Myc-His tag | Used for arabinose-inducible expression of Tme1 protein targeted to the periplasm in *E. coli* | (7) |
| $pTse1^{peri}$ | pPER5 plasmid | Used for arabinose- | This study |

| | | | |
|---|---|---|---|
| | containing the CDS of Tse1 (NP_250535.1) from *P. aeruginosa* PAO1 in frame with the N-terminal PelB signal peptide and the C-terminal Myc-His tag | inducible expression of Tse1 protein targeted to the periplasm in *E. coli* | |
| pBAD33.1$^{F}$ | pBAD33.1 with a FLAG tag inserted at the 3' end of the MCS | Used for arabinose-inducible expression of proteins | (7) |
| pHcp1b | pBAD33.1$^{F}$ plasmid containing the CDS of Hcp1b (WP_065788327.1) from *V. parahaemolyticus* strain CFSAN018764 in frame with the C-terminal FLAG tag of the plasmid | Used for arabinose-inducible expression of Hcp1b in *V. parahaemolyticus* | This study |
| pModule | pBAD33.1$^{F}$ containing the Hcp1b module (Hcp1b, DUF4225$^{18764}$, and Imm4225$^{18764}$) from *V. parahaemolyticus* strain CFSAN018764; Imm4225$^{18764}$ is cloned in-frame with the C-terminal Flag tag of the plasmid | Used for arabinose-inducible expression of the Hcp1b module in *V. parahaemolyticus* | This study |
| pPoNe | pBAD33.1$^{F}$ containing the *V. parahaemolyticus* 12-297/B VgrG1b auxiliary module with the effector and immunity pair PoNe/i$^{Vp\ 12\text{-}297/B}$ (*b5c30_rs14470-60*); PoNi$^{Vp\ 12\text{-}297/B}$ is cloned in-frame with the C-terminal FLAG tag of the plasmid | Used for arabinose-inducible expression of the PoNe/i$^{Vp\ 12\text{-}297/B}$ effector and immunity pair-containing VgrG1b module in *V. parahaemolyticus* | This study |
| pImm4225$^{18764}$ | pBAD33.1$^{F}$ plasmid containing the CDS of Imm4225$^{18764}$ in frame with the C-terminal FLAG tag of the | Used for arabinose-inducible expression of Imm4225$^{18764}$ in *E. coli* | This study |

| | | | |
|---|---|---|---|
| | plasmid | | |
| pDM4 | a *Cm*$^R$ and *oriV*$_{R6K}$-containing suicide vector | Used to generate deletions and substitutions in *Vibrio* genomes | (8) |
| pDM4:*hns* | pDM4 containing 1 kb upstream and 1 kb downstream of *vp1133* in its MCS | Used to delete *hns* in *V. parahaemolyticus* RIMD 2210633 | (9) |
| pDM4:*hcp1* | pDM4 containing 1 kb downstream and 1 kb upstream of *vp1393* in its MCS | Used to delete *hcp1* in *V. parahaemolyticus* RIMD 2210633 | (5) |
| pDM4:*vp1388* | pDM4 containing 1 kb upstream and 1 kb downstream of *vp1388* in its MCS | Used to delete *vp1388* in *V. parahaemolyticus* RIMD 2210633 | (10) |
| pDM4:*vpa1263* | pDM4 containing 1 kb upstream and 1 kb downstream of *vpa1263* in its MCS | Used to delete *vpa1263* in *V. parahaemolyticus* RIMD 2210633 | (10) |
| pDM4:*vp1415*$^{AAA}$ | pDM4 containing ~2.2 kb region encompassing 1.1 kb upstream and 1.1 kb downstream of the codons encoding histidines 563-4 in *V. parahaemolyticus* RIMD 2210633 *vp1415* in which these two codons were substituted to encode alanines | Used to substitute VP1415 histidines 563-4 for alanine in the *V. parahaemolyticus* RIMD 2210633 genome | (11) |
| pCLTR | *E. coli*-yeast-*Vibrio* shuttle vector; mobilizable; it contains Strep$^R$, Spec$^R$ and Cm$^R$ | Used to express proteins in *V. natriegens* | (11) |
| pImm | pCLTR containing the region encompassing *araC* to the rrnT1 terminator, amplified from the pImm4225$^{18764}$ plasmid | Used for arabinose-inducible expression of Imm4225$^{18764}$ in *V. natriegens* | This study |

### Supplementary Datasets

**Supplementary Dataset S1. *V. parahaemolyticus* genomes analyzed in this study.**

**Supplementary Dataset S2. Summary of T6SS gene clusters and auxiliary modules.**

**Supplementary Dataset S3. Analysis of *V. parahaemolyticus* T6SS gene clusters.**

**Supplementary Dataset S4. Analysis of *V. parahaemolyticus* T6SS auxiliary modules.**

**Supplementary Dataset S5. Similarity of *V. parahaemolyticus* Hcp proteins**

**Supplementary Dataset S6. List of DUF4225-containing proteins in bacterial genomes.**

### Supplementary Movies

**Supplementary Movie S1. Time-lapse microscopy.** *E. coli* cells containing a plasmid for arabinose-inducible expression, either empty or encoding DUF4225$^{18764\text{-Peri}}$ (Periplasmic DUF4225). A merge of phase contrast, RFP (detecting PI fluorescence), and GFP (detecting WGA-488 fluorescence) channels is shown.
